# Efficacy of Non-invasive Brain Stimulation on Vision: A Systematic Review and Meta-analysis

**DOI:** 10.1101/2022.11.22.517490

**Authors:** U.M. Bello, J. Wang, A.S.Y. Park, K.W.S. Tan, B.W.S. Cheung, B. Thompson, A.M.Y. Cheong

## Abstract

**Objective:** Multiple studies have explored the use of non-invasive brain stimulation (NIBS) to enhance visual function. These studies vary in sample size, outcome measures, and NIBS methodology. We conducted a systematic review and meta-analyses to assess the effects of NIBS on visual functions in human participants with normal vision.

**Methods:** We followed the PRISMA guidelines, and a review protocol was registered with PROSPERO before study commencement (CRD42021255882). We searched Embase, Medline, PsychInfo, PubMed, OpenGrey and Web of Science using relevant keywords. The search covered the period from 1^st^ January 2000 until 1^st^ September 2021. Comprehensive meta-analysis (CMA) software was used for quantitative analysis.

**Results:** Forty-nine studies were included, of which 19 were included in a meta-analysis (38.8%). Meta-analysis indicated acute (Hedges’s g=0.232, 95% CI: 0.023-0.442, *p*=0.029) and aftereffects (0.590, 95% CI: 0.182-0.998, *p*=0.005) of transcranial electrical stimulation (tES, including three different stimulation protocols) on contrast sensitivity. Visual evoked potential (VEP) amplitudes were significantly enhanced immediately after tES (0.383, 95% CI: 0.110-0.665, *p*=0.006). Both tES (0.563, 95% CI: 0.230 to 0.896, *p*=0.001)] and anodal-transcranial direct current stimulation (a-tDCS) alone (0.655, 95% CI: 0.273 to 1.038, *p*=0.001) reduced crowding in peripheral vision. The effects of NIBS on visual acuity, motion perception and reaction time were not statistically significant.

**Conclusions:** There are significant effects of visual cortex NIBS on contrast sensitivity, VEP amplitude, an index of cortical excitability, and crowding among normally sighted individuals. Future studies with robust experimental designs are needed to substantiate these findings in populations with vision loss.

**PROSPERO registration number:** CRD42021255882

**Highlights:**

- We conducted a meta-analysis and a systematic review on the efficacy of non-invasive brain stimulation for improving on visual function
- Visual cortex non-invasive brain stimulation can enhance contrast sensitivity, reduce crowding in peripheral vision and enhance visually evoked potential amplitude among normally sighted individuals.

## INTRODUCTION

Non-invasive brain stimulation (NIBS) enables the modulation of neural activity in targeted, superficial areas of the human brain. There are two primary NIBS techniques: transcranial magnetic stimulation (TMS) and transcranial electrical stimulation (tES).

TMS utilizes electromagnetic induction to generate brief electric currents within the stimulated brain area. This can be delivered as either single pulses or a string of repetitive pulses. Single pulses of TMS can generate action potentials that induce a motor or perceptual response. For example, TMS delivered to the primary motor cortex can cause peripheral muscle contraction (1) and TMS of the primary visual cortex can induce a phosphene percept (2). Repetitive pulses of TMS (rTMS) can increase or decrease cortical excitability within the stimulated brain region and alter the regional concentration of neurotransmitters such as gamma-aminobutyric acid (GABA) and glutamate (3). The nature of the rTMS effect on cortical excitability and local neurochemistry depends on the structure of the pulse train (4). Commonly used pulse trains include 1Hz and 10Hz stimulation frequencies as well as continuous and intermittent theta burst protocols (cTBS and iTBS) (5).

tES involves the delivery of an electrical current to the brain using head-mounted electrodes. tES stimulation protocols include transcranial direct current stimulation (tDCS), transcranial random noise stimulation (tRNS) and transcranial alternating current stimulation (tACS). tES does not induce action potentials but may alter membrane potentials (tDCS (6, 7), tRNS (8)), induce regional changes in neurotransmitter concentration (tDCS (9-11)), alter cortical excitability (tDCS (7, 12), tRNS (8, 13)), entrain patterns of neural activity (tACS (14)) and alter the signal to noise ratio within stimulated regions (tRNS, refer to Reed (15) for a review). NIBS has been used in multiple research contexts including the study of fundamental neurological processes, cognition (16-18), and the development of new therapeutic interventions (e.g. depression (19, 20), neurorehabilitation (21)).

Visual brain areas are attractive targets for NIBS research because regions such as the primary visual cortex and motion sensitive extrastriate area MT are close to the cortical surface with techniques such as visual psychophysics, electroencephalography, and magnetic resonance imaging available to measure the effects of the stimulation on neural activity and perception (22-24). In addition, NIBS is emerging as a promising tool for vision rehabilitation (25, 26). However, the literature on NIBS of visual brain areas is diverse with a wide range of different study designs, stimulation protocols, outcome measures and population samples. The aim of this structured review and meta-analysis was to assess whether visual cortex NIBS can enhance visual perception and/or modulate visual cortex activity (measured using visual evoked potentials). We did not include studies that used NIBS to induce “virtual lesions” or impair visual function to probe fundamental neurological processes. Our original plan was to review visual cortex NIBS studies involving either healthy or clinical populations (e.g. amblyopia (27, 28) or hemianopia (29, 30)). However, our literature search revealed that studies of clinical populations did not employ common study designs and were relatively few. We therefore limited our review to studies of healthy participants with normal vision.

## MATERIALS & METHODS

This systematic review conforms to the Preferred Reporting Items for Systematic Reviews and Meta-Analyses 2020 (PRISMA-2020) guidelines (31). We registered the review protocol with the International Prospective Register of Systematic Reviews (PROSPERO; Ref. No: <u>CRD42021255882</u>) in June 2021, prior to the initiation of the data extraction processes. We adopted the PICO (Participants, Intervention, Comparators and Outcome) format in generating the research question. The intervention was any form of NIBS [including tDCS, tACS, tRNS, and TMS], while the comparators included sham (placebo) NIBS. Outcomes of interest included psychophysical measures of contrast sensitivity, visual crowding, visual acuity, motion perception, visual evoked potentials (VEPs), and reaction time among others. The study conceptualization and development of the review protocol were undertaken by authors UMB, JW, BT and AMYC.

### Search strategy

A systematic search of PubMed, Embase, PsycINFO, Web of Science, Medline and OpenGrey databases was conducted from 1^st^ January 2000 until 1^st^ September 2021. The search terms were grouped under two themes, namely: ‘Brain area’, and ‘NIBS’. The electronic search involved combining terms under each theme using the Boolean operator ‘OR’. The search themes were then combined using the Boolean ‘AND’ (see Appendix 1 for details of the search themes/terms). Citation management software (EndNote X9, Clarivate Analytics, Philadelphia, Pennsylvania, USA) was used to organize the electronic search results and remove duplicates. Two of the authors (JW and UMB) independently conducted the electronic search. Any discrepancies during the independent search process were resolved by consulting a third author (AMYC). A thorough manual search of the reference lists of the identified studies and a forward reference search (via Google scholar) were also conducted.

### Study eligibility criteria

Studies were included if they: (i) assessed the effect of NIBS on enhancing visual functions among normally sighted individuals; (ii) included a sham stimulation control; (iii) were available in full text and (iv) written in English. We excluded studies that were: (i) conducted on individuals presenting with mental disorders, cognitive impairments, or visual impairments; (ii) used NIBS to disrupt or impair visual function, (iii) review protocols; (iv) systematic reviews; (v) conference abstracts and (vi) case studies.

### Article screening

The identified studies via electronic search processes were sequentially screened at the title, abstract and full text phases by two of the authors (JW and UMB). Any discrepancies identified by the two authors during the screening phases were resolved by discussion or consultation with the corresponding author (AMYC).

### Data extraction

The primary data for this study quantified the effect of NIBS on enhancing visual functions. Other relevant data extracted included the study reference, year of publication, title of study, study design, NIBS method, brain area stimulated, and visual function(s) measured. Data extraction was undertaken independently by JW and BWSC using an extraction tool designed in Microsoft Excel. Disagreements between the authors during the data extraction process were resolved by discussion or consultation with the other authors (BT and AMYC).

### Data analysis

Meta-analyses were conducted using the Comprehensive Meta-Analysis (CMA) software version 3.0 (Biostat Inc., Englewood, New Jersey, USA). Outcomes of studies that utilized protocols from the same NIBS delivery technique (tES or TMS) and reported findings on the same visual function were pooled for meta-analyses. Therefore tDCS, tRNS and tACS studies were pooled and rTMS and TBS studies were pooled. Similar studies with differing techniques for measuring a specific visual function could be pooled. Finally, studies with a common outcome measure were pooled. Examples include reaction time and VEP. Within each pooled group, we included all relevant studies and looked at *acute* (immediate, same day pre-vs. post-effects of NIBS), and *after* effects (same day, but at a designated time point after stimulation – i.e., 10 to 30 minutes post-stimulation). In the first instance, all related NIBS subtypes (tES or TMS) were combined for a general overview, and where there were enough studies, the stimulation protocol subtypes were analyzed separately. Study authors were contacted via email to obtain any missing data for the included studies. Unless otherwise indicated, stimulation was applied to the occipital lobe/primary visual cortex (V1). Data presented graphically were extracted using the GetData Graph Digitizer 2.26 (http://getdata-graph-digitizer.com/). Data reported as median and range were converted to mean and standard deviation (32). We adopted the bias-adjusted, standardized mean difference (SMD; Hedges’s g) to analyze the extracted data from the primary studies. The chi-square test (I^2^ statistics) was used to determine the degree of variance across studies (33), and a random-effects model was used for all the meta-analyses due to methodological heterogeneity among the studies. A p-value of <0.05 indicated statistical significance.

### Quality appraisals of the included studies

Two authors (ASYP, KWST) attempted to conduct quality ratings of the included studies using the Downs and Black quality rating tool (34), which consisted of 27 items. Ratings were conducted independently prior to comparison. However, it was noted that 14 randomly selected studies, were all rated of “poor” quality, suggesting that perhaps this instrument might not have been the most appropriate for the types of intervention studies included in this meta-analysis.

## RESULTS

### Characteristics of the included studies

In total, 5265 studies were identified through the electronic database and manual searches, among which 19 were included in the review after sequential screening of the title, abstract and the full text (27, 35-52). Appendix 2 summarizes the reasons for the studies excluded from meta-analysis.

The study flowchart detailing the search outcome and screening processes is presented in Figure Overall, the included studies recruited 674 participants. For the NIBS modalities adopted in the included studies, most studies utilized tDCS (n=14, 73.7%), then tRNS (n=3, 15.8%) and tACS (n=1, 5.3%). Another study utilized tRNS with tDCS (n=1, 5.3%). The visual functions examined among the studies were contrast sensitivity (n=7, 36.8%), reaction time (n=6, 31.6%), VEPs (n=4, 21.1%), motion perception (n=3, 15.8%), crowding (n=3, 15.8%), and visual acuity (n=2, 10.5%)^a^. Table 1 presents the study characteristics.

**Table 1.** Characteristics of the included studies (n=19)

| S/<br>No | Study<br>reference | Study<br>design | Outcome<br>measures | Age<br>(years) | Sex<br>(m:f) | N | NIBS | Stimu-<br>lation<br>site | Onlin-<br>e/<br>Offlin-<br>e | Mont-<br>age<br>(targe-<br>t-ref/ta-<br>rget) | Neu-<br>ro | Duratio-<br>n (min) | Stimu-<br>lation<br>sessio-<br>ns (n) | Intens-<br>ity<br>(mA /<br>MSO) | Size of<br>electrod-<br>e/coil<br>(cm<sup>2</sup>;<br>target-<br>referen-<br>ce) | Density<br>(mA/cm<sup>2</sup>) | Stimuli | Side-effect |
| --- | --- | --- | --- | --- | --- | --- | --- | --- | --- | --- | --- | --- | --- | --- | --- | --- | --- | --- |
| 1 | Barbieri<br>(2016)<br>(35) | between<br>subjects,<br>sham<br>controlled | face<br>perception<br>, object<br>perception<br>(RT) | 27 | 15:33 | 48 | atDCS | occipit-<br>o-<br>tempo-<br>ral<br>cortex | online<br>+<br>offline | PO8-FP1 | yes,<br>EEG | online:<br>24.6<br>sham:<br>24.6<br>offline:<br>20 | 1 | 1.5 | 25-25 | 0.08 | faces,<br>objects | no<br>side<br>effect |
| 2 | Battaglini<br>(2017)<br>(36) | within-subjects,<br>sham<br>controlled | motion<br>perception | / | 15:15 | 30 | atDCS<br>,<br>ctDCS | V5 | offline | left<br>V5/M-T-Cz | no | 12 | 4 | 1.5 | 5*7-5*7 | 0.043 | moving<br>dots | / |
| 3 | Battaglini<br>(2020)<br>(37) | within-subjects,<br>sham<br>controlled | CS | 25 ±<br>3.4 | 7:13 | 20 | tRNS | V1 | online | Oz-Cz | / | 15 | 4 | 1.5 | 7.2*6-11.5*9.5 | 0.03-0.01 | Gabor<br>patches | mild<br>skin<br>sensat-<br>ion |
| 4 | Behrens<br>(2017)<br>(38) | between-<br>subjects,<br>sham<br>controlled | CS | 24.5<br>±<br>3.5 | 12:12 | 24 | atDCS | V1 | offline | V1-Cz | yes,<br>MRI | 20 | 5 | 1.5 | / - 7*5 | 0.06 | Humphrey<br>perimetry<br>(central<br>10 °) | / |
| 5 | Bonder<br>(2018)<br>(39) | between-<br>subjects,<br>sham<br>controlled | VA (RT) | 26.3 | 8:22 | 30 | atDCS | occipit-<br>al<br>cortex | online | O1-FP2 | yes,<br>EEG | 15 | 1 | 1 | 5*5-7*5 | 0.04-0.03 | a black<br>square<br>with a gap<br>on either<br>side |  |
| 6 | Chen<br>(2021)<br>(40) | between<br>subjects,<br>sham<br>controlled | crowding | / | Exp 1:<br>23:22;<br>Exp 2:<br>21:24;<br>Exp 3:<br>16:12 | 118 | atDCS | occipit-<br>al<br>cortex | offline | P1/P2<br>-<br>ipsilat-<br>eral<br>cheek | yes,<br>EEG | 20 | 1 | 2 | 35-35 | 0.06 | gratings,<br>Sloan<br>letters | / |
| 7 | Contemori<br>(2019)<br>(41) | between-<br>subjects,<br>sham<br>controlled | crowding | 25 | 15:17 | 32 | tRNS<br>(100-640Hz<br>) | occipit-<br>al<br>cortex | online | Oz-Cz | no | 30 | 4 | 1.5 | 16-27 | 0.094 | single<br>white<br>letter,<br>crowded | / |

| S/<br>No | Study<br>reference | Study<br>design | Outcome<br>measures | Age<br>(years) | Sex<br>(m:f) | N | NIBS | Stimu-<br>lation<br>site | Online/<br>Offline | Mont-<br>age<br>(target/<br>target) | Neu-<br>ro | Duratio-<br>n (min) | Stimu-<br>lation<br>sessions (n) | Intens-<br>ity<br>(mA /<br>MSO) | Size of<br>electrode/coil<br>(cm<sup>2</sup>; target-reference) | Density<br>(mA/cm<sup>2</sup>) | Stimuli | Side-effect |
| --- | --- | --- | --- | --- | --- | --- | --- | --- | --- | --- | --- | --- | --- | --- | --- | --- | --- | --- |
|  |  |  |  |  |  |  |  |  |  |  |  |  |  |  |  |  | white<br>letters |  |
| 8 | +Ding<br>(2016)<br>(27) | within-subjects,<br>sham<br>controlled | CS, VEPs | 23 ±<br>2.3 | / | 27 | atDCS<br>,<br>ctDCS | occipit-<br>al<br>cortex | online<br>+<br>offline | Oz-Cz | yes,<br>EEG | 20 | 6 | 2 | 4*6-5*7 | 0.083-0.057 | Gabor<br>patches,<br>pattern-reversal<br>checkerbo-<br>ards | / |
| 9 | Dong<br>(2020)<br>(42) | within-subjects,<br>sham<br>controlled | EEG | 18-26 | 10:5 | 15 | atDCS | occipit-<br>al<br>cortex | offline | Oz-Cz | yes,<br>EEG | 21 | 2 | 2 | 35-35 | 0.06 | white<br>cross<br>fixation | no<br>side<br>effect |
| 10 | Fertonani<br>(2011)<br>(43) | between-<br>subjects,<br>sham<br>controlled<br>&<br>between-<br>subjects,<br>non-sham<br>controlled | orientatio-<br>n<br>discrimina-<br>tion (RT) | 21.7<br>±<br>2.5 | 42:42 | 84 | lf-<br>tRNS<br>( 0.1-100<br>Hz),<br>hf-<br>tRNS<br>(100-640<br>Hz),<br>atDCS<br>;<br>ctDCS | V1 | online | Oz-<br>right<br>arm | yes,<br>EEG | 22 | 1 | 1.5 | 16-60 | 0.025-0.06 | tilted<br>black lines | tDCS-induce<br>d<br>sensat-<br>ions<br>were<br>percei-<br>ved<br>strong-<br>er |
| 11 | He (2019)<br>(44) | within-subjects,<br>sham<br>controlled | CS | 23.4<br>±<br>1.9 | 16:11 | 27 | atDCS | occipit-<br>al<br>cortex | offline | Oz-Cz | yes,<br>EEG | 15 | 3 | 2 | 5*5-5*5 | 0.08 | gratings | no<br>side<br>effect |
| 12 | Kraft<br>(2010)<br>(45) | within<br>subjects,<br>sham<br>controlled | CS | 25.9<br>±<br>1.83 | 5:7 | 12 | atDCS<br>,<br>ctDCS | occipit-<br>al<br>cortex | offline | O1/O2<br>-Cz | yes,<br>MRI | 15 | 3 | 1 | 5*5-<br>7*10 | 0.04-0.014 | Humphrey<br>perimetry | / |
| 13 | Lau<br>(2021)<br>(46) | within-subjects,<br>sham<br>controlled | VEPs | 28.7 | 6:14 | 20 | atDCS<br>,<br>ctDCS | V1 | offline | Oz-Cz | yes,<br>EEG | 20 | 3 | 2 | 5*7-5*7 | 0.06 | pattern-reversal<br>checkerbo-<br>ard | / |

| S/ No | Study reference | Study design | Outcome measures | Age (years) | Sex (m:f) | N | NIBS | Stimulation site | Online/ Offline | Montage (target-ref/target) | Neuro | Duration (min) | Stimulation sessions (n) | Intensity (mA / MSO) | Size of electrode/coil (cm<sup>2</sup>; target-reference) | Density (mA/cm<sup>2</sup>) | Stimuli | Side-effect |
| --- | --- | --- | --- | --- | --- | --- | --- | --- | --- | --- | --- | --- | --- | --- | --- | --- | --- | --- |
| 14 | Nakazono (2020) (47) | Exp 1: single arm<br>Exp 2&3: within-subjects, sham-controlled | CS, EEG, VEPs | 26.8 ± 7.8 | Exp 1: 6:7<br>Exp 2: 10:7<br>Exp 3: 7:8 | Exp 1: 13<br>Exp 2: 17<br>3:<br>Exp 3: 15 | tACS (10, 20 Hz) | V1 | offline | Oz-Cz | yes, EEG | 20 | Exp 1: 2<br>Exp 2&3: 3 | 1 | 3.5*3.5-7*5 | 0.08-0.03 | Pattern-reversal checkerboard, reversing black and white fields, Gabor patches | itching, flickering |
| 15 | Raveendran (2020) (48) | within-subjects, sham controlled | crowding | / | / | 13 | atDCS | V1 | online | Oz-Cz | yes, EEG | 20 | 2 | 2 | 5*5-5*5 | 0.08 | Gabor patches with flankers | / |
| 16 | Reinhart (2016) (49) | Exp 1: single arm<br>Exp 2-5: within subjects, sham controlled | CS, VA, VEPs (RT) | Exp 1: 22.0 ±0.9<br>Exp 2: 25.3 ±1.3<br>Exp 3: 23.1 ±1.2<br>;<br>Exp 4: 22.4 ±1.3 | Exp 1: 9:11<br>Exp 2: 13:7<br>Exp 3: 7:13<br>Exp 4: 10:10<br>Exp 5: 14:6 | 20 | atDCS , ctDCS | occipitoparietal cortex | offline | Exp 1: P1/P2-left/right cheek (ipsilateral)<br>Exp 2: left/right cheek (ipsilateral)-P1/P2<br>Exp 3: C3/C4 - left/right | yes, EEG | 20 | Exp 1: 3<br>Exp 2-5: 2 | Exp 1: 1/1.5/2<br>Exp 2-5: sham/2. | 19.25-52 | 0.1-0.038 | Vernier stimulus, Snellen letters, gratings | tingling, itching |

| S/<br>No | Study<br>reference | Study<br>design | Outcome<br>measures | Age<br>(years) | Sex<br>(m:f) | N | NIBS | Stimu-<br>lation<br>site | Online/<br>Offline | Mont-<br>age<br>(target-ref/target) | Neu-<br>ro | Duratio-<br>n (min) | Stimu-<br>lation<br>sessions (n) | Intens-<br>ity<br>(mA /<br>MSO) | Size of<br>electrode/coil<br>(cm<sup>2</sup>; target-reference) | Density<br>(mA/cm<sup>2</sup>) | Stimuli | Side-effect |
| --- | --- | --- | --- | --- | --- | --- | --- | --- | --- | --- | --- | --- | --- | --- | --- | --- | --- | --- |
|  |  |  |  | ;<br>Exp<br>5:<br>20.4<br>±1.8 |  |  |  |  |  | ht<br>cheek<br>(ipsila-<br>teral)<br>Exp 4-5:<br>P1/P2-left/rig-<br>ht<br>cheek<br>(ipsila-<br>teral) |  |  |  |  |  |  |  |  |
| 17 | van<br>Koningsb-<br>ruggen<br>(2016)<br>(50) | between<br>subjects,<br>sham<br>controlled | attentional<br>capture<br>(RT) | 23.8<br>±<br>3.6 | 20:40 | 60 | tRNS<br>(100-640<br>Hz) | Latera-<br>l<br>occipit-<br>al<br>cortex | online | PO7-PO8 | yes,<br>EEG | 20 | 1 | 1 | 5*7-5*7 | 0.03 | black lines<br>in empty<br>coloured<br>circles | / |
| 18 | Wu<br>(2020)<br>(51) | between<br>subjects,<br>sham<br>controlled | motion<br>perception | / | 28:0 | 28 | atDCS | V5 | offline | V5-Cz | no | 20 | 1 | 1.5 | 5*7-5*7 | 0.043 | moving<br>white dots | / |
| 19 | Zito<br>(2015)<br>(52) | within-subjects,<br>sham<br>controlled | motion<br>perception<br>(RT),<br>shape<br>perception | 30.5<br>±<br>5.1 | 11:10 | 21 | aHD-tDCS,<br>cHD-tDCS | V5 | offline | P4,<br>OZ,<br>TP8,<br>&<br>PO10-PO8 | yes,<br>EEG | 20 | 3 | 2 | / | / | moving<br>dots,<br>ellipses | no<br>effect |

**Abbreviations:** + Those studies included participants with amblyopia and healthy controlled (but only healthy controlled were considered in this meta-analysis); N = Sample size
Outcome measures: CS (contrast sensitivity); RT (reaction time); VA (visual acuity); VEPs (visual evoked potentials)
Types of stimulation:
1) Transcranial electrical stimulation (tES): atDCS (anode transcranial direct current stimulation); ctDCS (cathode transcranial direct current stimulation); aHD-tDCS (anodal high-definition transcranial direct current stimulation); cHD-tDCS (cathode high-definition transcranial direct
stimulation); tACS (transcranial alternating current stimulation); tRNS (transcranial random noise stimulation); lf-tRNS (low-frequency transcranial random noise stimulation); hf-tRNS (high-frequency transcranial random noise stimulation) 2) Transcranial magnetic stimulation (TMS): rTMS (repetitive transcranial magnetic stimulation); cTBS (continuous theta burst stimulation).
Application of neuro-navigation (neuro): EEG (electroencephalogram); MRI (magnetic resonance imaging).
Site of stimulation: Cz (central zero); FP (frontal pole); MT (middle temporal visual area); Oz (occipital zero); PO (posterior occipital); TP (temporal pole); V1 (primary visual cortex).

### Quantitative analysis of whether NIBS can enhance visual function

#### 1) Contrast sensitivity

##### 1.1 Acute effect of tES (a-tDCS, tRNS, and tACS) on contrast sensitivity

Included studies measured the same-day effects of a single NIBS session on contrast sensitvity. The pooled analysis involved seven studies, among which five utilized a-tDCS (27, 38, 44, 45, 49), one adopted tACS (47) and one utilized tRNS (37). In studies that measured multiple outcomes, only the contrast sensitivity results were included (27, 47, 49). Contrast sensitivity was measured using perimetry (38, 45), a 10 cycles per degree (cpd) Gabor patch (27), or stimuli presented at a range of spatial frequencies (37, 44, 47). If the study measured contrast sensitivity at more than one spatial frequency, the results for the highest spatial frequency was chosen, because the most challenging condition was expected to show the greatest NIBS-induced enhancement. Spatial frequencies selected included 9 cpd (47), 10 cpd (27), and both 7 cpd and 12 cpd for the study by Battaglini (37) because the authors explicitly hypothesized that sensitivity for both higher spatial frequencies would be enhanced by the tRNS. For studies that measured contrast sensitivity at more than one retinal eccentricity, the measures for central vision were selected for meta-analysis to provide consistency across studies. Nakazono et al. (47) compared alpha and beta tACS to a sham condition. Both stimulation frequencies were included in the meta-analysis. Similarly, Battaglini et al. (37) used vertical and 45-degree oriented Gabors for their contrast detection tasks. Both orientations were included in the meta-analysis. We pooled the effect of active stimulation against sham conditions for the analysis (Figure 2). The result indicated a statistically significant acute effect of tES stimulation (Hedges’s g=0.232, 95% CI: 0.023-0.442, *p*=0.029) on contrast sensitivity in normally sighted participants.

**Figure 1:**
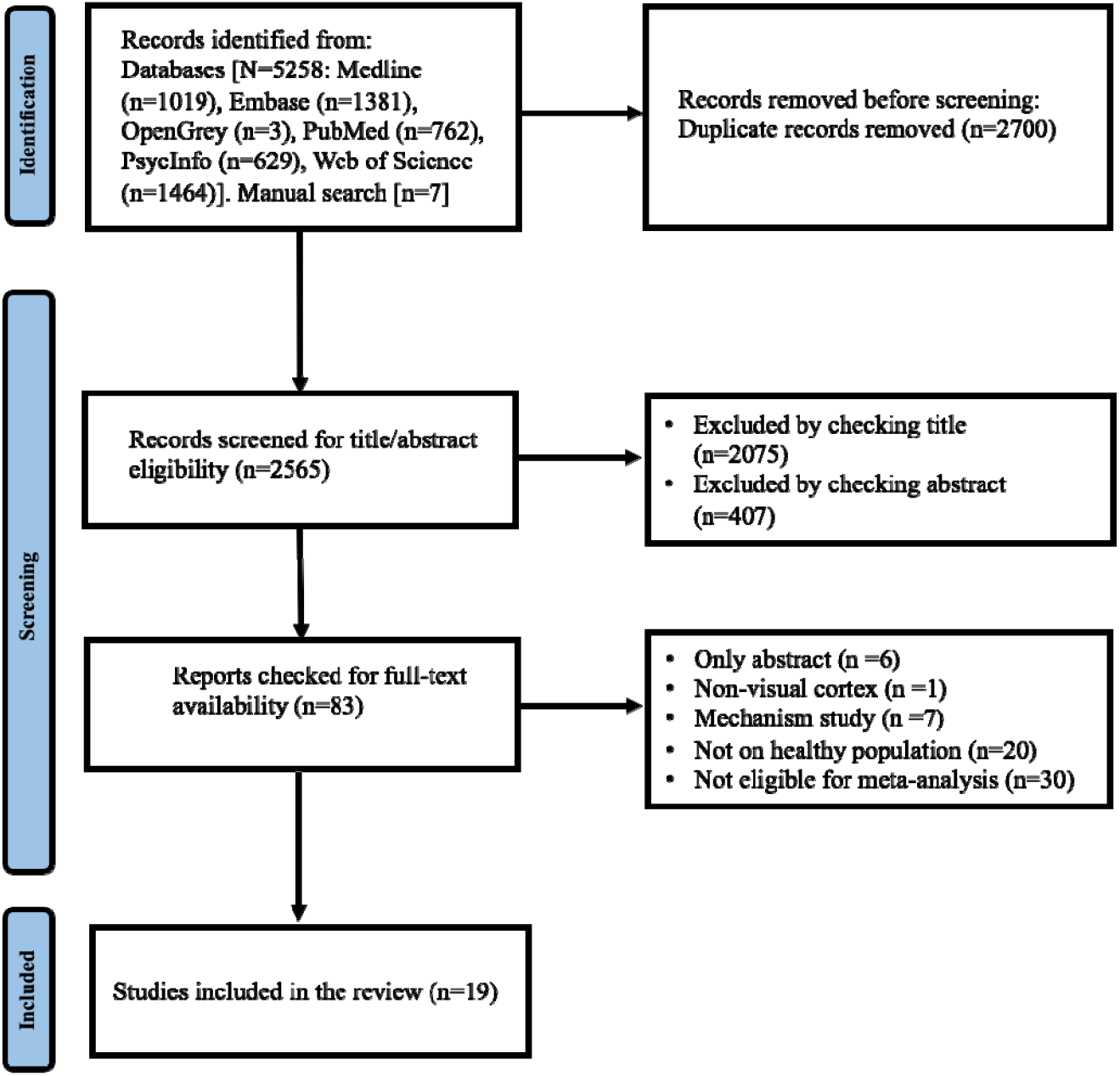
Study flowchart

**Figure 2:**
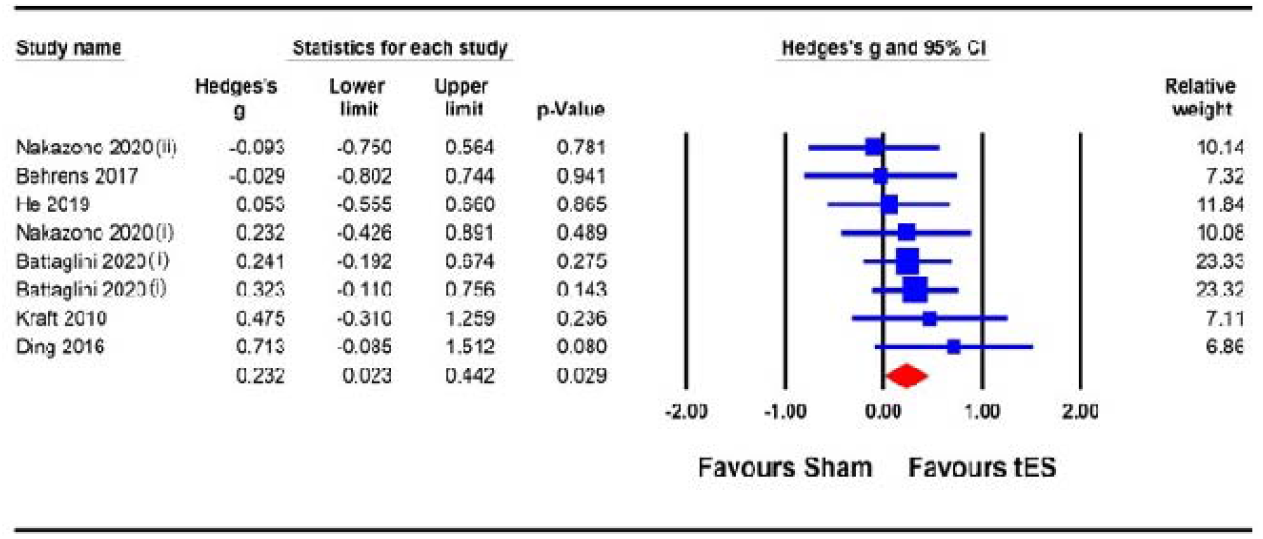
Acute effect of tES (a-tDCS, tRNS, and tACS) on contrast sensitivity. ***Description:*** Meta-analyses for Nakazono (2020) were separated for data on 9.0 cpd alpha (acute) and 9.0 cpd beta (acute), represented as Nakazono 2020(i) and Nakazono 2020(ii); Ding 2016, acute effect; Meta analyses for Battaglini (2020) were separated for data using contrast stimuli of 45 degree and vertical, represented as Battaglini 2020(i) and Battaglini 2020(ii).

##### 1.2 Acute effect of anodal tDCS (a-tDCS) on contrast sensitivity

A single session acute effect of a-tDCS on contrast sensitivity is illustrated in Figure 3. Of those studies included in section 1.1, the a-tDCs studies were pooled for the analysis (27, 38, 44, 45). There was a trend favouring an effect of a-tDCS stimulation on contrast sensitivity as per the main analysis, but this failed to reach statistical significance (Hedges’s g=0.262, 95% CI: -0.101-0.625, *p*=0.158).

**Figure 3:**
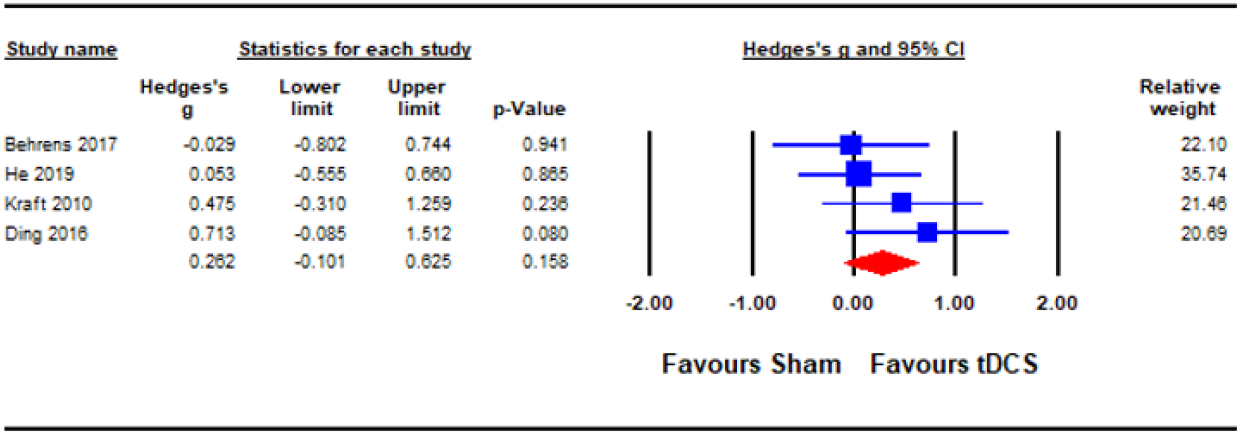
Acute effect of a-tDCS on contrast sensitivity. ***Description:*** Ding 2016, acute effect.

##### 1.3 Aftereffect of tES (a-tDCS and tACS) on contrast sensitivity

Nakazono (47) and Ding (27) reported aftereffects of tES on contrast sensitivity measured 10- and 30-minutes post-stimulation respectively (Figure 4). A meta-analysis revealed a statistically significant aftereffect of tES stimulation on contrast sensitivity (Hedges’s g=0.590, 95% CI: 0.182-0.998, *p*=0.005).

**Figure 4:**
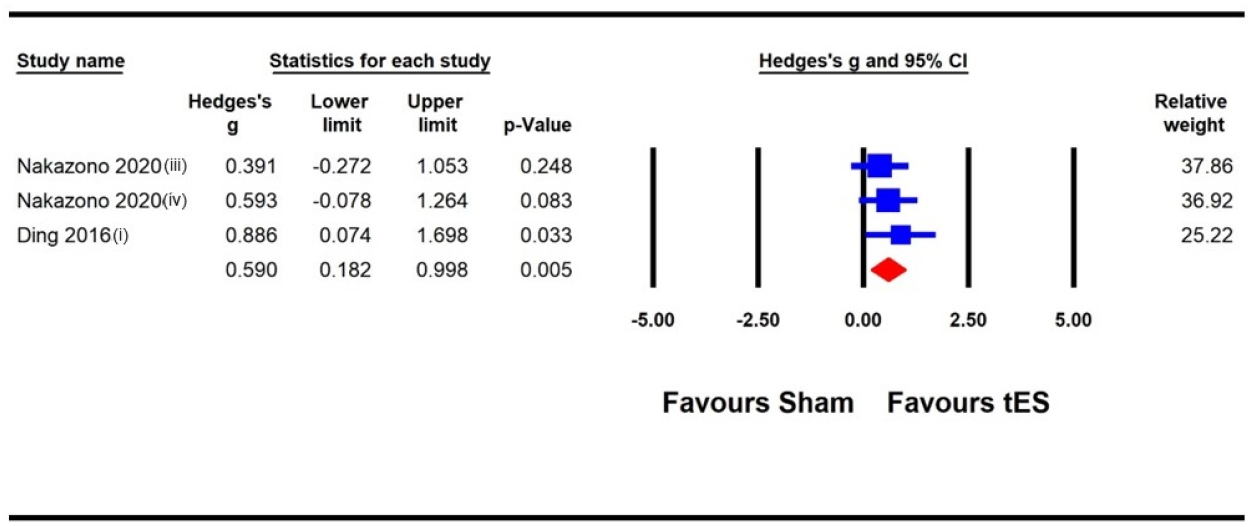
Aftereffect of tES (a-tDCS and tACS) on contrast sensitivity. ***Description:*** Meta-analyses for Nakazono (2020) were separated for data on 9.0 cpd beta (at 10 mins post) and 9.0 cpd alpha (at 10 mins post), represented as Nakazono 2020(iii) and Nakazono 2020(iv); Ding 2016(i), 30 mins post.

#### 2) Visual evoked potentials (VEPs)

##### 2.1 Acute effect of tES (a-tDCS and tACS) on VEP amplitude

Four studies were pooled for analysis to assess the acute effect of tES on VEP amplitude, three utilized a-tDCS (27, 42, 46) and one adopted tACS (47). In studies that measured the acute effects of NIBS on different visual functions (e.g., contrast sensitivity and VEPs), the results from the VEP measure were taken (27, 47). Different components of VEPs were estimated in the studies, including amplitude of P100-N75 (27, 47), amplitude of the alpha activity over the parieto-occipital area (42), and N1 and P1 amplitudes (46) (both included in the analysis). Similarly, where alpha and beta tACS were utilized in a study (47), the effect of each stimulation condition against a sham effect was extracted for the analysis. We pooled the effect of active stimulation against sham condititons for the analysis (Figure 5). The result indicated a statistically significant increase in VEP amplitude imediately after tES at visual cortex (Hedges’s g=0.383, 95% CI: 0.110-0.655, *p*=0.006).

**Figure 5:**
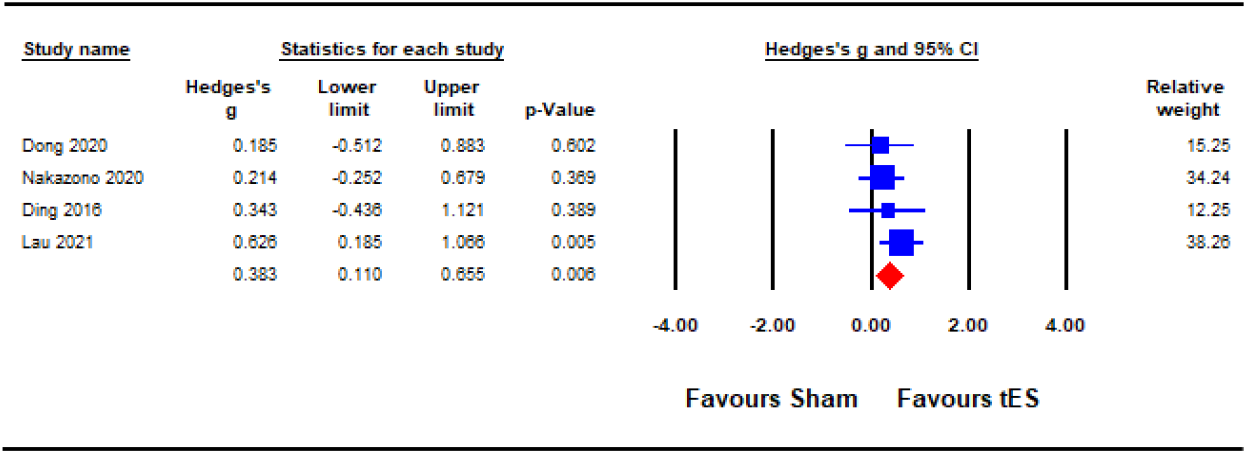
Acute effect of tES (a-tDCS and tACS) on visual evoked potentials (VEPs). **Description:** Nakazono 2020, combined effect of alpha and beta tACS; Lau 2021, combined effect of N1 and P1 amplitudes.

#### 3) Crowding

##### 3.1 Acute effect of tES (a-tDCS and tRNS) on crowding

Three studies were pooled for analysis to assess the acute effect of tES on crowding, two utilized a-tDCS (40, 48), and one adopted tRNS (41). In the study with multiple experiments involving different groups of participants (40), data from each experiment were pooled separately in th analysis. The results for NIBS applied to the hemisphere contralateral to the presented stimuli against a sham condition were pooled in the analysis (40). The earliest effect of a-tDCS on crowding (5 minutes post-stimulation) reported by Raveendran and colleagues (48) was pooled for analysis. The analysis (Figure 6) indicated a statistically significant effect of tES on crowding (Hedges’s g=0.563, 95% CI: 0.230 to 0.896, *p*=0.001).

**Figure 6:**
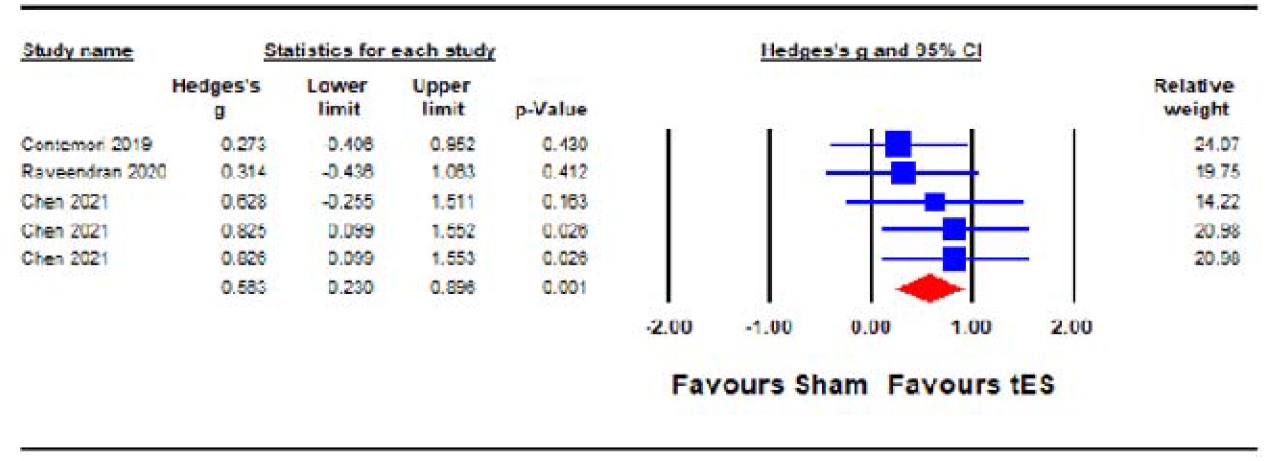
Acute effect of tES (a-tDCS and tRNS) on crowding. **Description:** Chen 2021 (each line represents the outcome of experiments 1-3).

##### 3.2 Acute effect of a-tDCS on crowding

To assess the acute effect of a-tDCS on crowding (independent of tRNS), data from the two studies that used a-tDCS (40, 48) were pooled for analysis. The result (Figure 7) indicated a statistically significant effect of a-tDCS on crowding (Hedges’s g=0.655, 95% CI: 0.273 to 1.038, *p*=0.001).

**Figure 7:**
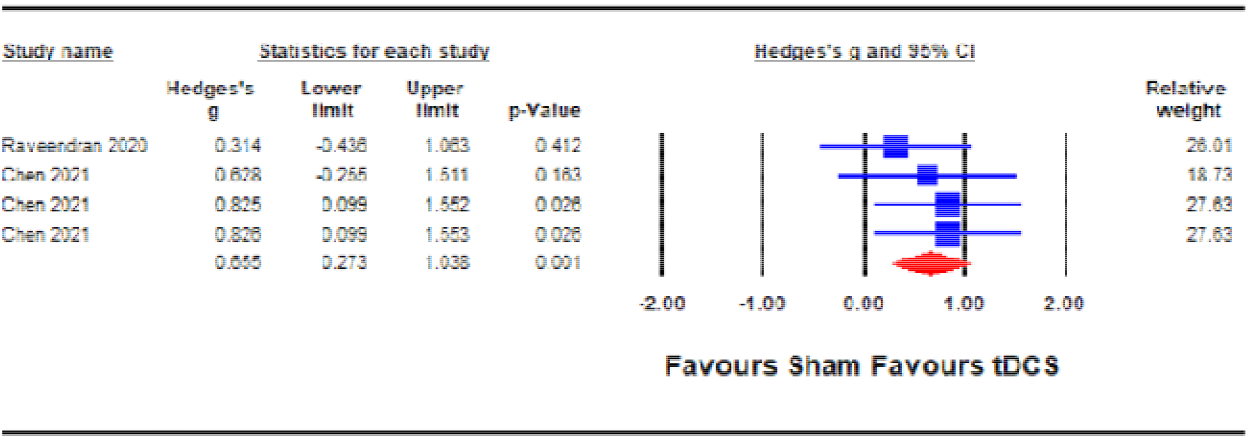
Acute effect of a-tDCS on crowding. **Description:** Chen 2021 (each line represents the outcome of experiments 1-3).

#### 4) Visual acuity

##### 4.1 Acute effect of a-tDCS on visual acuity

To assess the acute effect of a-tDCS on visual acuity, two studies (39, 49) were pooled for analysis. In a study that measured the acute effects of a-tDCS on different visual functions (visual acuity, contrast sensitivity and VEPs), the results from the visual acuity measure were taken (49). The pooled effect for the active stimulation condition in each study was compared to sham conditions (Figure 8). The result indicated a statistically non-significant effect of a-tDCS on visual acuity (Hedges’s g=0.408, 95% CI: -0.056 to 0.872, *p*=0.085).

**Figure 8:**
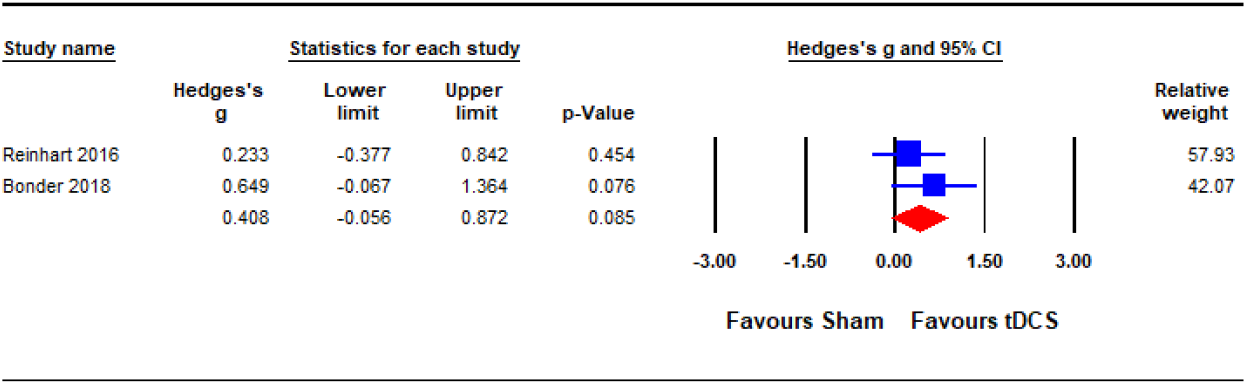
Acute effect of a-tDCS on visual acuity.

#### 5) Motion perception

##### 5.1 Acute effect of a-tDCS on motion perception

The pooled analysis to assess the acute effect of a-tDCS on motion perception involved three studies (36, 51, 52). All studies stimulated extrastriate cortical area V5 (middle temporal area; MT). In the study that measured the acute effects of a-tDCS on different visual functions (motion and shape perception), the results from the motion perception measure were taken (52). The pooled effect for each of the study were compared against sham control conditions (Figure 9). The result indicated a statistically non-significant effect of a-tDCS on motion perception (Hedges’s g=0.802, 95% CI: -0.458 to 2.063, *p*=0.212).

**Figure 9:**
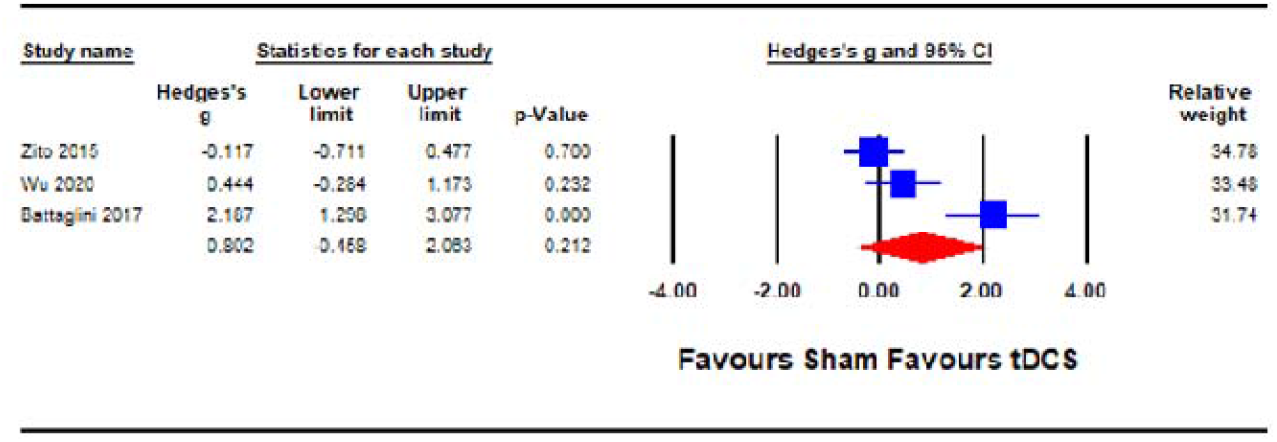
Acute effect of a-tDCS on motion perception.

#### 6) Reaction time

##### 6.1 Acute effect of tES (tRNS and a-tDCS) on reaction time

Reaction time was analysed as a proxy of vision-related cognitive processing. Six studies were pooled for the analysis to assess the acute effect of tES on reaction time, five utilized a-tDCS (35, 39, 49, 52) and two adopted tRNS (43, 50). In studies that measured the acute effects of NIBS on different visual functions (e.g., face/object/motion perception, visual acuity, VEPs, contrast sensitivity, attentional capture and reaction time), the results from the reaction time measure were taken (35, 39, 43, 49, 50, 52). When a study reported the effect of multiple NIBS protocols on same group of participants (for example tRNS and a-tDCS) (43), both effects were pooled in the tES meta-analysis (Figure 10), respectively. Similarly, in a study with dual experimental/stimulation conditions (face and object recognition) (35), the effects of both conditions on the reaction time were combined and pooled in the analysis. Outcome of the analysis illustrated a statistically non-significant effect of tES on reaction time (vision-related cognitive processing) (Hedges’s g=1.001, 95% CI: -0.405 to 2.406, *p*=0.163).

**Figure 10:**
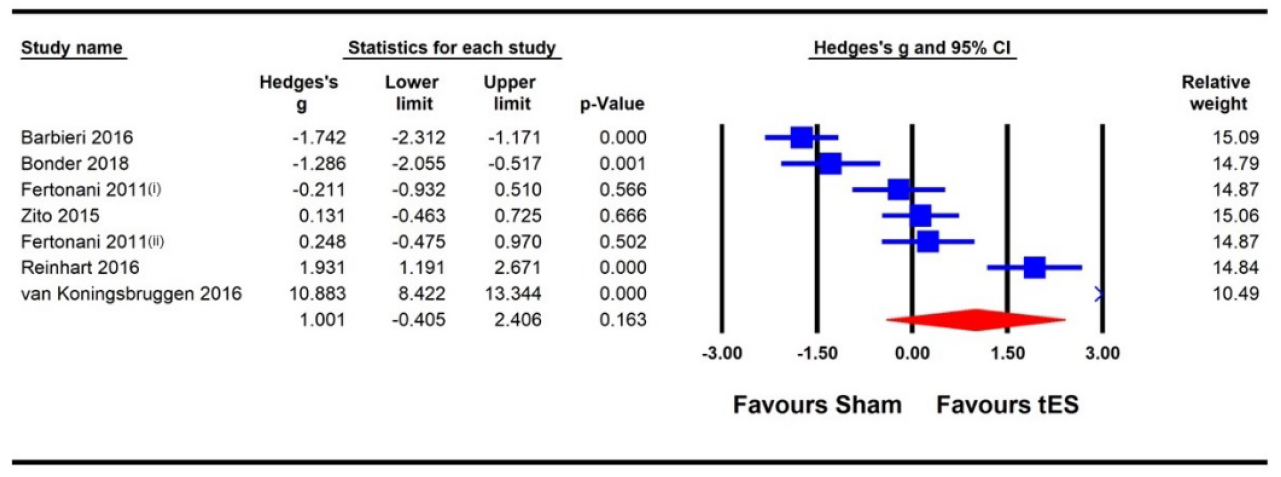
Acute effect of tES (a-tDCS and tRNS) on reaction time. ***Description:*** Barbieri 2016 (included data for combined effect of a-tDCS face and object tasks); Meta-analysis for Fertonani (2011) were separated for data on a-tDCS and tRNS, represented as Fertonani 2011(i) and Fertonani 2011(ii).

##### 6.2 Acute effect of a-tDCS on reaction time

To assess the acute effect of a-tDCS on reaction time, data from the five studies that used a-tDCS in the sub-section 6.1 (35, 39, 43, 49, 52) were pooled. The result (Figure 11) indicated a statistically non-significant effect of a-tDCS on reaction time (Hedges’s g=-0.241, 95% CI: - 1.474 to 0.991, *p*=0.701).

**Figure 11:**
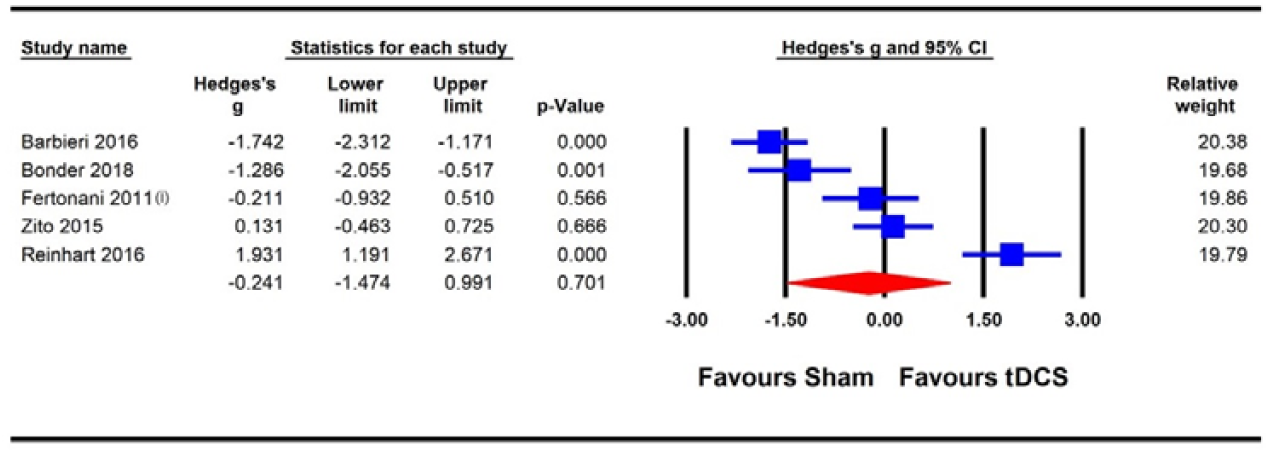
Acute effect of a-tDCS on reaction time. **Description:** Barbieri 2016, combined effect of a-tDCS face and object tasks; Fertonani 2011(i), effect of a-tDCS.

## DISCUSSION

To recapitulate, the aim of this structured review and meta-analysis was to assess whether visual cortex NIBS could enhance visual perception and/or modulate visual cortex activity. Studies exploring the use of NIBS to enhance normal vision have employed a diverse range of experimental designs with time scales that range from the acute effects of a single NIBS session to multi-session studies that combine NIBS with perceptual learning. This diversity resulted in limited opportunities for meta-analyses. However, by pooling across different tES stimulation protocols and differing methodologies for assessing a common outcome measure, we were able to assess the effects of a single tES session vs. sham stimulation on contrast sensitivity, VEP amplitude, visual crowding, visual acuity, motion perception, and reaction time.

Both contrast sensitivity (27, 37, 38, 44, 45, 47, 49) and visual crowding (40, 41, 48) were significantly enhanced by visual cortex tES relative to sham within our meta-analyses, and were examined at different timescales relative to stimulation. Results from our meta-analyses showed beneficial acute effects of tES in enhancing contrast sensitivity (Figure 2) and reduced crowding (Figure 6). The sub-analysis of studies that only employed a-tDCS revealed improvements in crowding following stimulation (Figure 7), but not for contrast sensitivity (Figure 3). Our meta-analyses looking at later time points (i.e., aftereffects) could only be performed for contrast sensitivity as there was only one study investigating the effect of NIBS on crowding. We observed that tES was effective in modulating contrast sensitivity at a fixed time point after stimulation (Figure 4), indicating that the effects of stimulation on improving contrast sensitivity persisted beyond the stimulation period. Despite only one study measuring the aftereffects of a-tDCS on crowding (48) (in this case lateral-inhibition, a low-level mechanism that may contribute to crowding), the study reported a larger effect at 30 minutes vs. 5 minutes after stimulation, which is in line with the time-scale of the effects observed in the contrast sensitivity meta-analysis.

From a mechanistic perspective, the meta-analysis of changes in VEP amplitude (27, 42, 46, 47) following visual cortex tES vs. sham revealed enhanced cortical excitability (i.e., larger VEP amplitudes) following tES (Figure 5). Increased cortical excitability may enhance neural sensitivity to contrast and weaken lateral inhibition mechanisms that contribute to crowding. The connection between tES and increased cortical excitability may be mediated by the relative concentration of the inhibitory neurotransmitter GABA and the excitatory neurotransmitter glutamate within the stimulated area. Reduced GABA concentration within motor cortex following a-tDCS has been reported by multiple studies (53, 54) and it is possible that tES may have a similar effect when applied to the visual cortex. Within this framework, the delayed effects of tES on contrast sensitivity (27, 47) (and perhaps crowding (48)) could reflect a gradual change in GABA concentration that continues for a period after the stimulation session. However, the time course of tES effects on GABA concentration remains unclear and it is also unknown whether the effects of tES on GABA concentration are the same for the motor and visual cortices. It is worth noting that indirect evidence exists suggesting that visual cortex tES does not influence GABA (55). Therefore, while the effects of visual cortex tES on contrast sensitivity, crowding, and VEP amplitude are supported by our meta-analyses, the underlying mechanisms require investigation.

Meta-analyses revealed no evidence for the effectiveness of tES on visual acuity, motion perception or reaction time. The visual acuity and motion perception meta-analyses included the fewest individual experiments (2 for visual acuity (39, 49) and 3 for motion perception (36, 51, 52)) with considerable variations in the visual stimuli used to measure the outcomes. The small sample combined with significant protocol differences may have limited our power to detect an effect. It is possible that measures of visual acuity and motion perception differ from contrast sensitivity and crowding in their response to tES. For motion perception, it is also possible that area MT responds to tES in a way that is distinct from that of the primary visual cortex. However, additional studies are required to fully address these questions.

The reaction time meta-analysis included 7 experiments and revealed high variability across studies with two reporting longer reaction times following tES (35, 39), three reporting no effect (43, 52) and two reporting shorter reaction times (49, 50); one with a moderate effect size (49) and the other with a Hedge’s G greater than 5 (50). Reaction times can be affected by multiple variables including attention, task complexity, participant instructions, and speed accuracy trade-off. The studies included in the reaction time meta-analysis differed considerably in the types of visual stimuli and tasks employed and therefore it is perhaps not surprising that tES of cortical regions responsible for early, low-level visual processing did not produce consistent effects across studies.

The diverse nature of the NIBS and vision literature forced us to pool across different tES protocols, visual stimuli, and experimental designs in our meta-analyses. Therefore, our results should be interpreted with caution. In particular, a non-significant meta-analysis may reflect important variations in experimental parameters rather than no effect of the stimulation itself. In addition, our inclusion of multiple independent experiments from a single publication may have amplified study-specific sources of bias. As the literature on NIBS and vision continues to develop, future meta-analyses may be able to adopt more stringent analyses criteria.

## CONCLUSION

Meta-analyses revealed evidence for the effectiveness of visual cortex tES compared to sham stimulation on modulating contrast sensitivity and crowding. These effects were accompanied by evidence for a significant increase in visual cortex excitability indexed by VEP amplitude following tES. Despite the diversity of study designs in the current NIBS and vision literature, the results of this review indicate that NIBS can enhance at least some visual functions and strengthen the foundation for the application of NIBS in studies of vision rehabilitation.

## Appendix 1. Search strategy

Appendix 1: Search strategy
| Search Terms | Search Themes |  |
| --- | --- | --- |
|  | *Brain area* | *Noninvasive brain stimulation (NIBS)* |
| Search Terms | “Primary visual cortex” | “Transcranial direct current stimulation” |
|  | “V1” | “tDCS” |
|  | “Extrastriate cortex” | “Transcranial alternating current stimulation” |
|  | “V2” | “tACS” |
|  | “V3” | “Transcranial random noise stimulation” |
|  | “V3a” | “tRNS” |
|  | “V4” | “Transcranial magnetic stimulation” |
|  | “V5” | “TMS” |
|  | “Middle temporal” | “Repetitive transcranial magnetic stimulation” |
|  | “MT” | “rTMS” |
|  | “Medial superior temporal” | “Continuous theta burst stimulation” |
|  | “MST” | “cTBS” |
|  | “Occipital pole” | “Theta burst stimulation” |
|  | “Occipital lobe” | “TBS” |

## Appendix 2. Reasons for the studies excluded from the meta-analysis

Appendix 2: Reasons for the studies excluded from the meta-analysis
| S/N | Study | Visual functions | Reason(s) |
| --- | --- | --- | --- |
| 1 | Abuleil 2021 (55) | Binocular rivalry, mixed percept duration | The only paper in category (visual function) |
| 2 | Antal 2004 (56) | Visuo-motor coordination | No sham data |
| 3 | Antal 2004 (57) | VEP | No sham group |
| 4 | Antal 2004 (58) | Visuo-motor coordination | The only paper in category (visual function) |
| 5 | Battaglini 2020 (59) | Visual temporal resolution | The only paper in category (visual function) |
| 6 | Battaglini 2020 (60) | Motion perception | No appropriate data for meta-analysis (only a sensitivity parameter - d'prime was provided and no raw data were provided by the corresponding author) |
| 7 | Bocci 2018 (61) | VA and VEP | No sham group |
| 8 | Cabral-Calderin 2015 (62) | Bistable perception | The only paper in category (visual function) |
| 9 | Campana 2016 (63) | Visual motion adaptation | The only paper in category (visual function) |
| 10 | Chaieb 2008 (64) | VEP, CS | No appropriate data for meta-analysis (only percentage change were reported, and no raw data were provided by the corresponding author) |
| 11 | Costa 2015 (65) | Size and distance of judgement | The only paper in category (visual function) |
| 12 | Costa 2015 (66) | Visual field | The only paper in category (visual function) |
| 13 | Costa 2015 (67) | VEP | Sweep VEP instead of pattern/ flash VEP was used. Parameters reported were very different from other studies (amplitude of suprathreshold) |
| 14 | Costa 2012 (68) | Color discrimination | The only paper in category (visual function) |
| 15 | Hanson 2015 (69) | Overlay and surround suppression | No sham group |
| 16 | Heinrichs-Graham 2017 (70) | Oscillatory activity | The only paper in category (visual function) |
| 17 | Kim 2019 (71) | VEP (steady-state) | No appropriate data for meta-analysis (signal-to-noise ratio and no raw data were provided by the corresponding author) |
| 18 | Laczo 2012 (72) | CS, visual attention | No appropriate data for meta-analysis (only percentage change were reported and no raw data were provided by the corresponding author) |
| 19 | Larcombe 2019 (73) | Motion perception | Combined tDCS with visual training was reported (no data for pure tDCS effect) |
| 20 | Peters 2013 (74) | Contrast sensitivity, VEP | No data for the acute effect (i.e. immediately after stimulation) |
| 21 | Pirulli 2013 (75) | Orientation and discrimination | The only paper in category (visual function) |
| 22 | Ranieri 2019 (76) | VEP after rTMS | The only paper in category (TMS stimulation) |
| 23 | Richard 2015 (77) | CS | No sham group |
| 24 | Schaeffner 2019 (78) | Contrast polarity and depth perception | The only paper in category (visual function) |
| 25 | Sczesny-Kaiser 2016 (79) | VEP | VEP parameters reported were very different from other papers (amp ratio (A2s/A1)) |
| 26 | Somer 2020 (80) | RT | No appropriate data for meta-analysis (no pre-post data, no sham data and no raw data were provided by the corresponding author) |
| 27 | Spiegel 2012 (81) | Surround suppression | The only paper in category (visual function) |
| 28 | Thompson 2008 (26) | CS after rTMS | The only paper in category (stimulation) |
| 29 | Van Meel 2016 (82) | Object recognition, orientation judgement | The only paper in category (visual function) |
| 30 | Waterston 2010 (83) | CS (course orientation and discrimination) at peripheral vision | The only paper in category (visual function) |

## Footnotes

a Note that some studies included more than one visual functions. Hence, the sum of the total percentage exceeds 100%.

